# Reaction time sensitivity to spectrotemporal modulations of sound

**DOI:** 10.1101/2022.01.13.476175

**Authors:** Lidwien C.E. Veugen, A. John van Opstal, Marc M. van Wanrooij

## Abstract

We tested whether sensitivity to acoustic spectrotemporal modulations can be observed from reaction times for normal-hearing and impaired-hearing conditions. In a manual reaction-time task, normal-hearing listeners had to detect the onset of a ripple (with density between 0-8 cycles/octave and a fixed modulation depth of 50%), that moved up or down the log-frequency axis at constant velocity (between 0-64 Hz), in an otherwise-unmodulated broadband white-noise. Spectral and temporal modulations elicited band-pass filtered sensitivity characteristics, with fastest detection rates around 1 cycle/oct and 32 Hz for normal-hearing conditions. These results closely resemble data from other studies that typically used the modulation-depth threshold as a sensitivity criterion. To simulate hearing-impairment, stimuli were processed with a 6-channel cochlear-implant vocoder, and a hearing-aid simulation that introduced separate spectral smearing and low-pass filtering. Reaction times were always much slower compared to normal hearing, especially for the highest spectral densities. Binaural performance was predicted well by the benchmark race model of binaural independence, which models statistical facilitation of independent monaural channels. For the impaired-hearing simulations this implied a “best-of-both-worlds” principle in which the listeners relied on the hearing-aid ear to detect spectral modulations, and on the cochlear-implant ear for temporal-modulation detection. Although singular-value decomposition indicated that the joint spectrotemporal sensitivity matrix could be largely reconstructed from independent temporal and spectral sensitivity functions, in line with time-spectrum separability, a substantial inseparable spectral-temporal interaction was present in all hearing conditions. These results suggest that the reaction-time task yields a valid and effective objective measure of acoustic spectrotemporal-modulation sensitivity.

## Introduction

Human speech and other complex sounds in the natural environment are typically dynamic signals that rapidly change in amplitude over both time and frequency. Fluctuations in the temporal domain provide information about the rhythm of speech, such as syllable and word boundaries, whereas variations in the spectral domain are essential for formant and voice pitch perception (Liberman 1996). Sensitivity to these joint spectral and temporal modulations is deemed crucial for the identification of complex sound features (McDermott and Simoncelli 2011) and for speech comprehension (Elliott and Theunissen 2009; Shannon et al. 1995).

Spectrotemporal dynamic ripples have been introduced in psychoacoustics to investigate the spectrotemporal modulation sensitivity of auditory perception. Ripples are broadband noise stimuli that are modulated sinusoidally in amplitude over time and/or frequency (Bernstein and Green 1998; Supin et al. 1994). Ripples are ideal to assess hearing performance as they represent features of, but are not recognizable as, naturalistic sounds. Sensitivity of the healthy human auditory system has been studied thoroughly with ripples and generally shows a band- or low-pass response to spectral and temporal modulations, reflecting the limits of auditory sensitivity at higher modulation rates (Chi et al. 1999; Viemeister 1998; Zheng, Escabí, and Litovsky 2017). Speech understanding is thought to relate mostly to joint spectrotemporal sensitivity. Chi et al. (1999) reported that in normal-hearing listeners the modulation transfer function of combined spectrotemporal ripples is highly separable, as it can be well approximated by the product of a single temporal and spectral filter. Separability implies that the joint spectrotemporal sensitivity can be directly obtained from pure temporal and spectral sensitivity measurements.

In the present study we used manual reaction times to construct the spectrotemporal modulation transfer function (stMTF), rather than the conventionally used modulation detection or discrimination thresholds. Research in monkeys shows that reaction times systematically depend on acoustic modulation rates (Massoudi et al. 2014). Several models have been proposed to explain the underlying process of response latency in reaction-time tasks (Ratcliff and Van Dongen 2011). It is commonly assumed that a decision signal rises with accumulating evidence of the stimulus, until a certain threshold is reached that triggers the response. As such, reaction times are directly related to the difficulty of a task and could thus provide more detailed information on the audibility of spectrotemporal ripples.

Furthermore, reaction times allow for testing the presence or absence of binaural integration based on monaural responses, by comparing binaural reaction times against the prediction of a so-called ‘race model’. In such a model, the signals from either ear compete independently to reach the detection threshold, so that the response latency is determined by the winner of an independent parallel race between the two ears (Raab 1962). Due to statistical facilitation, this race to threshold leads to faster reaction times for binaural stimulation than for monaural stimuli, as the distribution of minimum monaural reaction times yields faster responses than those produced by either ear (Gielen, Schmidt, and Van Den Heuvel 1983; Hershenson 1962). However, when this so-called redundant stimulus effect differs from the race-model prediction, it could imply true binaural integration in an underlying neural interaction process (Gielen et al. 1983; Miller 1982; Schröter, Ulrich, and Miller 2007).

We tested whether reaction times are an objective measure having convergent validity of auditory sensitivity to moving ripples with various spectrotemporal modulations for normal-hearing listeners. We assessed the extent of separability of joint spectrotemporal sensitivity and investigated how binaural listening affected modulation sensitivity compared to monaural listening conditions by comparing the data with the race-model prediction. As a validation of our reaction-time paradigm, we also collected data under more challenging impaired-hearing simulations that are known to affect temporal and spectral sensitivity (Bacon and Viemeister 1985; Golub et al. 2012; Henry, Turner, and Behrens 2005; Moore and Glasberg 2001) and binaural integration (Ausili et al. 2019; Sharma et al. 2019, 2021; Lidwien C.E. Veugen et al. 2016).

## Methods

### Listeners

Six listeners participated in this study (3 male, ages 20-25 years), none of whom reported a history of auditory deficits. All listeners had normal hearing (< 20 dB HL) in both ears from 125 to 8000 Hz. Except for two of the authors, listeners were naïve to the purpose of the experiments. We included the data from the two authors as they were highly similar to the data from the naïve listeners (see Supplemental Materials) and excluding those data did not appreciably affect the results and conclusions. The study was approved by the Local Ethics Committee of the Radboud University Nijmegen (protocol number 40327.091.12).

### Apparatus

Listeners were seated in an acoustically shielded sound chamber. Stimuli were presented through TDH 39 headphones (Telephonics Corporation, Farmingdale, NY, USA). For sound processing and data acquisition we used Tucker Davis Technologies System 3 (Alachua, FL, USA). Stored sounds were sent via the PC to a real-time processor (RP2.1) at a sampling rate of 48828.125 Hz, and passed through a programmable attenuator (PA5). Stimuli were set at a comfortable, well-audible loudness of 65 dB(A) (calibrated using a KEMAR head calibration set, connected to a Brüel & Kjaer measuring amplifier type 2610 [Nærum, Denmark]).

### Stimuli

Dynamic ripples were created in MATLAB [version R2012a; Mathworks Inc., Natick, MA, USA] as described by Depireux, Simon, Klein, and Shamma (2001). The carrier of these stimuli consisted of a broadband spectrum of multiple harmonic tones, each described by:

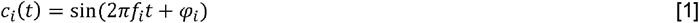

where *t* is time (s), *f*_*i*_ frequency (Hz) of the *i*-th harmonic, and *φ*_*i*_ is its phase (rad). In this experiment, the broadband carrier consisted of 128 harmonic tones, equally spaced (20 tones/octave) over 6.4 octaves (250 Hz-20.4 kHz). All components had random phase except for the first (*φ*_0_ =0). The *i*-th frequency was determined by *f*_*i*_ = *f*_0_ 2^*i*/20^ with *f*_0_ = 250 Hz the lowest frequency, and *i* = 0 - 127. All harmonic tones had the same amplitude, effectively yielding the same spectrum as white noise.

The spectrotemporal envelope determined the ripple fluctuations in amplitude over time and/or frequency:

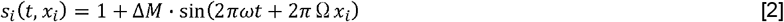

with *t* is time (s), *x*_*i*_ is the position on the frequency axis (in octaves above the lowest frequency), Δ*M* is the modulation depth, ω is the ripple velocity (Hz) and Ω is ripple density (cycles/octave). Unpublished data on freefield, normal-hearing ripple detection from our lab suggested that the actual value of the modulation depth is not very crucial. Therefore, we set the modulation depth rather arbitrarily to 0.5 for all components. Testing only one modulation depth reduced the number of potential parameter combinations and trials. This decreased the duration of an experimental session, which was already substantial.

Together the carrier and the modulator formed the dynamic ripple in our experiments as follows:

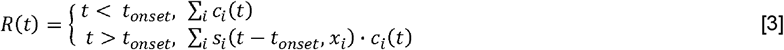

The modulated sounds were thus preceded by a non-modulated harmonic complex (*c*_*i*_(*t*)) with a randomized duration (*t*_*onset*_) between 700 and 1200 ms with a step size of 100 ms. Moving ripples were presented for a duration of maximally 3 s, with velocities of 0 Hz and ±[0.5; 1; 2; 4; 8; 16; 32; 64] Hz and densities [0; 0.25; 0.5; 0.75; 1; 2; 4; 8] cycles/octave, yielding a total of (17 velocities x 8 densities =) 136 different stimuli.

### Cochlear implant simulation

cochlear-implant vocoder simulations were created using a previously described method by Litvak et al. (Litvak et al. 2007) that models the Advanced Bionics Harmony cochlear-implant processor. Briefly, the vocoder algorithm works as follows. After resampling the input signal to 17.4 kHz, the vocoder applied a high-pass pre-emphasis filter (cut-off at 1.5 kHz). Then, the signal was band-pass filtered by a short-time Fourier transform with 256 bins and 75% temporal overlap (192 bins). Bins were grouped into 6 non-overlapping logarithmically spaced channels (Fig. 1B; at center frequencies: 452, 715, 1132, 1792, 2836 and 4488 Hz). Random-phase noise bands with similar center frequencies were modulated with amplitudes equal to the square root of the total energy in the channel. The channels were summed, and inverse short-time Fourier transformed to reproduce a temporal waveform for presentation to the listeners.

**Fig. 1.**
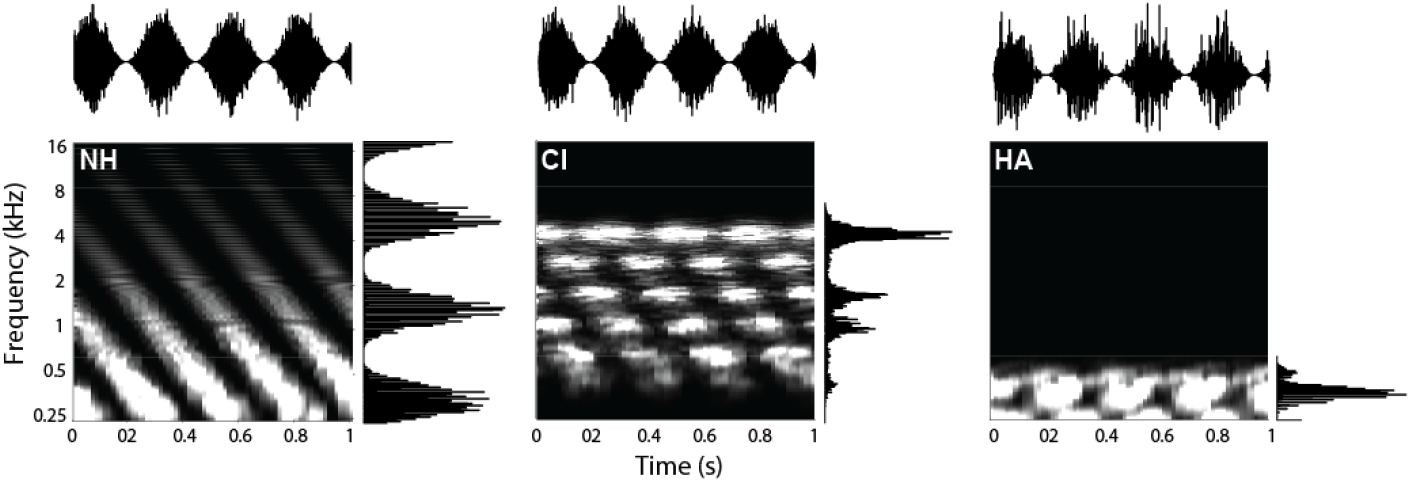
Moving-ripple spectrograms. Ripple with velocity 4 Hz and density 0.5 cycles/octave) for the normal-hearing condition (NH), cochlear implant simulation (CI) and hearing-aid simulation (HA). Examples of the temporal and spectral modulations of this ripple presented separately are shown at the top and right of each panel. The signals on the top row represent the temporal waveforms for a purely amplitude-modulated sound (4 Hz, 0 cycles/octave). The signal on the right of each panel visualizes a pure stationary spectral ripple modulation (0 Hz, 0.5 cycles/octave). For clarity, the sound is shown after t_onset_ (at t=0).

We used 6 vocoder channels to simulate hearing via a cochlear implant, as cochlear-implant users are typically unable to effectively utilize information from all available cochlear-implant channels (Henry and Turner 2003). Normal-hearing listeners have shown similar performance as cochlear-implant users for speech understanding scores in quiet with 4-6 channels (Loizou, Dorman, and Tu 1999). This is in line with pilot experiments in our lab that revealed that five normal-hearing listeners achieved a performance level of ∼80% in a consonant-vowel-consonant recognition test, when the words were vocoded with only 6 channels.

### Hearing aid simulation

hearing-aid simulations were generated by using a fourth-order Butterworth low-pass filter with a cut-off at 500 Hz, mimicking residual hearing in the lower frequencies present in the bimodal cochlear implant users of our previous studies (bimodal here refers to listeners using a cochlear implant in one ear, and a hearing aid in the other; Sharma et al. 2019; Lidwien C. E. Veugen et al. 2016). Additionally, the loss of frequency selectivity (spectral smearing) was simulated as previously described by Baer and Moore (1994). Asymmetrically broadened auditory filters were used with broadening factors 6 and 3 for the lower and upper branch respectively, as these are representative for moderate-to-severe hearing impairment (Glasberg and Moore 1986). The cochlear implant and hearing-aid simulated stimuli were normalized to the same root-mean-squared value as the original non-vocoded sounds (Fig. 1 visualizes the effect of the cochlear implant and hearing-aid simulation on ripples).

### Paradigm

Listeners were instructed to press a button as quickly as possible when they heard the sound change from static noise to modulated ripple. Modulated ripples lasted for 3000 ms, unless the button was pressed, in which case the sound was ended prematurely, and the next trial was initiated after a brief (0.5 - 1 s) period of silence between each trial. If the button was pressed before ripple onset, the trial was reiterated, but no more than 4 times. The outcome measure of the experiment was the listener’s manual reaction time, which was defined as the time between the onset of the ripple and the moment the button was pressed.

We tested five different listening conditions; acoustic stimuli were presented 1) monaurally (monaural normal hearing), 2) monaurally via cochlear-implant vocoder (unimodal cochlear implant), 3) monaurally via hearing-aid simulation (unimodal hearing aid), 4) binaurally (binaural normal hearing), 5) binaurally via bimodal cochlear implant and hearing-aid simulation (bimodal). In monaural conditions, both ears were tested separately. In the bimodal condition, cochlear implant and hearing aid were tested in both the right and left ear in different sessions. We did not test the binaural unimodal listening conditions (cochlear implant-cochlear implant or hearing aid-hearing aid). Each stimulus was presented 5 times in each listening condition. A complete data set thus contained a total of 6120 stimuli, which were split in 12 sessions of 30-40 minutes, each containing 510 trials. Sessions were distributed over 6 days of two sessions each. Ripples and conditions were presented in pseudo-randomized order.

Because of time constraints, data collection was not fully completed in the four naïve listeners. Two naïve listeners completed 11 out of 12 sessions, with all ripples measured at least twice. The other two listeners completed 9 out of 12 sessions (20 and 23 ripples not measured, respectively; 83 and 141 ripples were presented only once in these listeners). The four naïve listeners performed one training session under normal hearing conditions prior to the recording sessions, to become familiarized with the ripple stimuli and experimental procedures. We observed no systematic change in the average reaction times during the training session for these four listeners, or over the time course of the experimental sessions for all six listeners (e.g., for binaural trials, the mean reaction time decreased marginally by -6 ms [95% confidence interval lower, upper bound: -42, +47 ms] in the 401^st^ to 500^th^ trial compared to the first 100 trials, yielding a P value of a two-sided permutation t-test of 0.815; changes in mean reaction times varied between -27 to +21 ms across listening conditions, all of which did not reach significance [P>0.323]). This observation indicates that procedural learning effects did not confound the reaction-time data.

### Analysis

Data analysis was performed with custom-written MATLAB software. Reaction times generally show a skewed distribution with an extended tail towards longer reaction times. To obtain normally distributed data (Carpenter, Reddi, and Anderson 2009), the reaction-time data were transformed to their reciprocal (1/reaction time), referred to here as ‘promptness’ (1/s). This also allows the measurements to be more readily interpreted as sensitivity measures to the different spectrotemporal modulations, as a higher/lower promptness (as opposed to a shorter/longer reaction time) indicates a higher/lower sensitivity. Responses were pooled across listeners and ears for grand average analyses. Reaction times below 150 ms (clear anticipatory responses) were removed from the analysis. If a response was not initiated within 3 s (considered a sign for inattentiveness, or of an inability to detect the ripple), we set the response time (promptness) to 3 s (1/3 s^-1^). Non-responses were found in 10% of the trials under normal-hearing conditions and in 46% of the trials of the hearing-impaired conditions (especially at the high spectral modulations). We do not explicitly account for the percentage of non-responses but note that in our analyses a higher number of non-responses would yield a median promptness (reaction time) closer to 1/3 s^-1^ (3 s). The non-modulated sound (velocity 0 Hz and density 0 cycles/octave) served as a catch stimulus, to determine the guess (or false-alarm) rate of the participant. The guess rate varied from 7% for binaural normal hearing, to 21% for monaural hearing-aid listening, with the average guess rate across conditions at 12%.

### Spectrotemporal transfer function

For each of the five listening conditions (monaural and binaural normal-hearing, and monaural and bimodal cochlear implant and hearing-aid simulation), we calculated the mean promptness per ripple to construct a two-dimensional spectrotemporal modulation transfer function *stMTF*(*ω*,Ω) as a joint function of ripple density, Ω, and ripple velocity, ω. Similarly, we determined the temporal modulation transfer function tMTF, F(*ω*), and the spectral modulation transfer function sMTF, G(Ω), for the 0-density and 0-velocity stimuli, respectively.

### Separability

To analyze the degree of separability of the stMTF, we applied singular value decomposition (SVD) for all listening conditions. SVD transforms the stMTF into two unitary matrices containing temporal and spectral singular vectors, respectively, and a rectangular diagonal matrix that contains the singular values: stMTF(ω,Ω)=F(ω)·Σ·G(Ω). In case of a fully separable stMTF, the spectral and temporal components are independent of each other and the total of all 136 spectrotemporal responses can be expressed by the vectorial outer product of a single temporal F_1_(ω) (17 components) and spectral G_1_(Ω) (8 components) modulation transfer function, as follows:

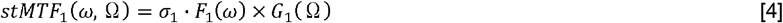

with σ_*1*_ the largest singular value.

We calculated the separability index (cf. inseparability index, as used by, for example, Massoudi et al. 2015; Versnel, Zwiers, and van Opstal 2009), which ranges between zero (totally inseparable) to one (fully separable), and is based on the relative dominance of the first SVD component:

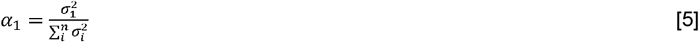

The separable stMTF estimate was reconstructed according to Eq. 4.

We also reconstructed the stMTF based on the first two singular values, according to

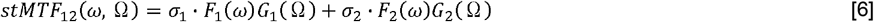

and determined the relative contribution of the first two SVD components:

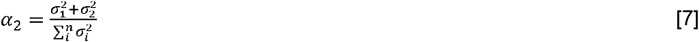

Before applying the SVD, the stMTF data was centered by subtracting the mean promptness for each listening condition. This mean was added to the reconstructions.

### Race model

We compared the observed reaction times for binaural hearing with the quantitative predictions of performance based on the monaural reaction times, using the race model of statistical facilitation. This model assumes independence of the two monaural processes (Gielen et al. 1983; Raab 1962). Any violation to the race model suggests neural interactions when processing the input from both ears:

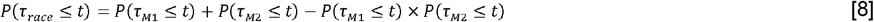

with P(τ ≤ t) the cumulative probability function (CDF) of an observed reaction time τ at time t; M_1_ and M_2_ represent monaurally-presented stimuli (normal hearing, cochlear implant and hearing aid). We estimated the cumulative probability density functions (CDF) from the promptness values. The race model CDF was constructed from the two monaural CDFs according to Eqn. 8. For comparative purposes, we calculated the difference in the medians (at the 50% cumulative probability level) between actual performance and race-model predictions. Ripples, for which fewer than 10 responses were collected, were discarded from this analysis because no reliable CDF could be constructed. Non-responses (reaction times > 2500 ms) were also discarded from the race-model analysis.

### Statistics

Data were always reported as mean values ± 1 standard deviation. We also calculated 95% confidence intervals of promptness for the pure temporal and spectral ripples. As a criterion of significance for a statistical difference we took p<0.05.

## Results

### Reaction-time task

We will first illustrate the systematic dependence of the manual reaction times on the acoustic conditions with the data of one listener (Fig. 2). Pure amplitude-modulated noises elicit cumulative distributions of reaction times in the binaural listening condition that shift systematically with the velocity of the stimulus (Fig. 2A). The cumulative distributions are plotted on a probit scale as a function of the reciprocal of reaction times (Carpenter and Williams 1995; Corneil et al. 2002). In this format, the data points for each velocity fall closely on a straight line indicating that the promptness responses form a normal distribution. One may note that the singly-most distinguishing feature of these lines is that they are shifted versions of each other, with the slopes being similar across the velocity modulations. This may suggest that the mean reaction time and its inverse, promptness (the promptness at 50% cumulative probability) is a good point estimate of the effect of a ripple’s velocity on reaction speed. Indeed, the mean promptness of this listener systematically and monotonically increases when velocity is increased (Fig. 2D). Also, for spectral modulations, reaction time distributions (Fig. 2B) and mean promptness (Fig. 2E) vary systematically with a ripple’s density, albeit that reaction speed seems to decrease with increasing density. Listening condition also affected this listener’s response speed, as exemplified for a [1 Hz, 0 cycles/octave] modulation (Figs. 2C, F), with binaural hearing eliciting the fastest responses and the monaural hearing-aid simulation yielding the slowest responses. In the following sections, we will quantify this relationship with promptness and the ripple modulation parameters for all listeners.

**Fig. 2.**
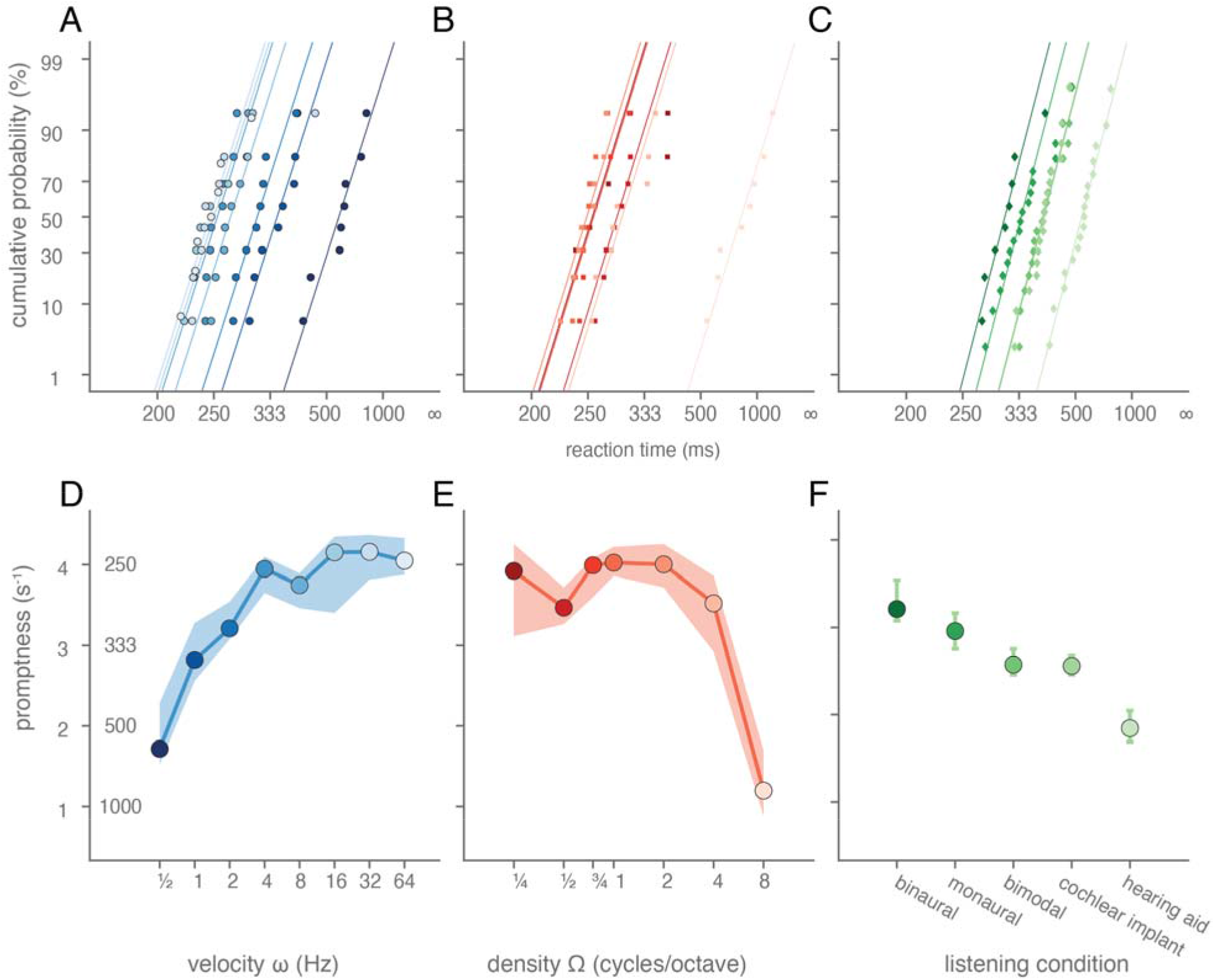
Example reaction times to temporally and spectrally modulated target sounds. A-C) Reaction time distributions plotted in the form of cumulative percentage probability, on a probit scale, as a function of reciprocal reaction time (promptness), measured in one listener (L1) in various listening regimes. The temporal and spectral modulation of the target sound and the listening condition was varied, leading to changes in the listener’s detection speed. Each curve corresponds to a different modulation or listening condition. A) Reaction time distributions to amplitude modulations (density = 0 cycles/octave) in a binaural listening condition are shown. As each distribution seemed to lie close to a straight line, regression lines are drawn, for illustrative purposes. Reaction time distributions to B) spectral modulations (velocity = 0 Hz) in a binaural listening condition and to C) different listening conditions for a single amplitude-modulated target sound (velocity = 2 Hz, density=0 cycles/octave) are shown. D-F) Mean promptness (cf. reaction time, see left ordinate) is plotted as a function of D) velocity, E) density, and F) listening condition (stimulus parameters are the same as in A-C). Cochlear implant and hearing aid indicate cochlear-implant and hearing-aid simulation, respectively. Colours in each plot (2A-F) indicate velocity, density and listening condition as indicated by the abscissa-values of the coloured, filled circles (in 2D-F). Coloured margins of error around point estimates indicate 95% confidence intervals of the mean.

### Temporal-only modulations

We will first elaborate on how reaction times reflect the detection of temporal-only modulations (Fig. 3A). For the normal-hearing listening conditions (binaural and monaural), the mean promptness as a function of velocity for the purely temporal amplitude modulations (density = 0 cycles/octave) resembled a high-pass characteristic (Fig. 3A, dark blue circles and light blue triangles). Responses were fastest (higher promptness) for the highest absolute velocities and were slower for lower velocities. If the sounds were vocoded, simulating hearing device processing (cochlear implant, hearing aid, bimodal), the promptness dropped especially for the higher velocities, so that the curve exhibited bandpass properties. Responses were now fastest for intermediate absolute velocities and were slower for both higher and lower velocities. Overall, both the maximal promptness and at which velocity this was attained were affected by listening condition; the fastest responses, with an average promptness of 3.3 (monaural, light blue triangles), 3.5 (binaural, dark blue circles), 2.5 (bimodal, pink squares), 2.5 (cochlear implant, light green diamonds) and 2 s^-1^ (hearing aid, dark green triangles) were observed at ±32, ±32, ±16, ±16, ±8 Hz, respectively. The longer response times to pure amplitude modulations under impaired-hearing conditions clearly implicate increased difficulty in the detection of temporal modulations. Differences in response times between binaural and monaural listening will be considered in more detail below with race-model predictions. Responses to upward (<0 Hz) and downward (>0 Hz) moving ripples were very similar; correlation coefficients between the responses to up- and downward ripples were between 0.91-0.99 for all listening conditions.

**Fig. 3.**
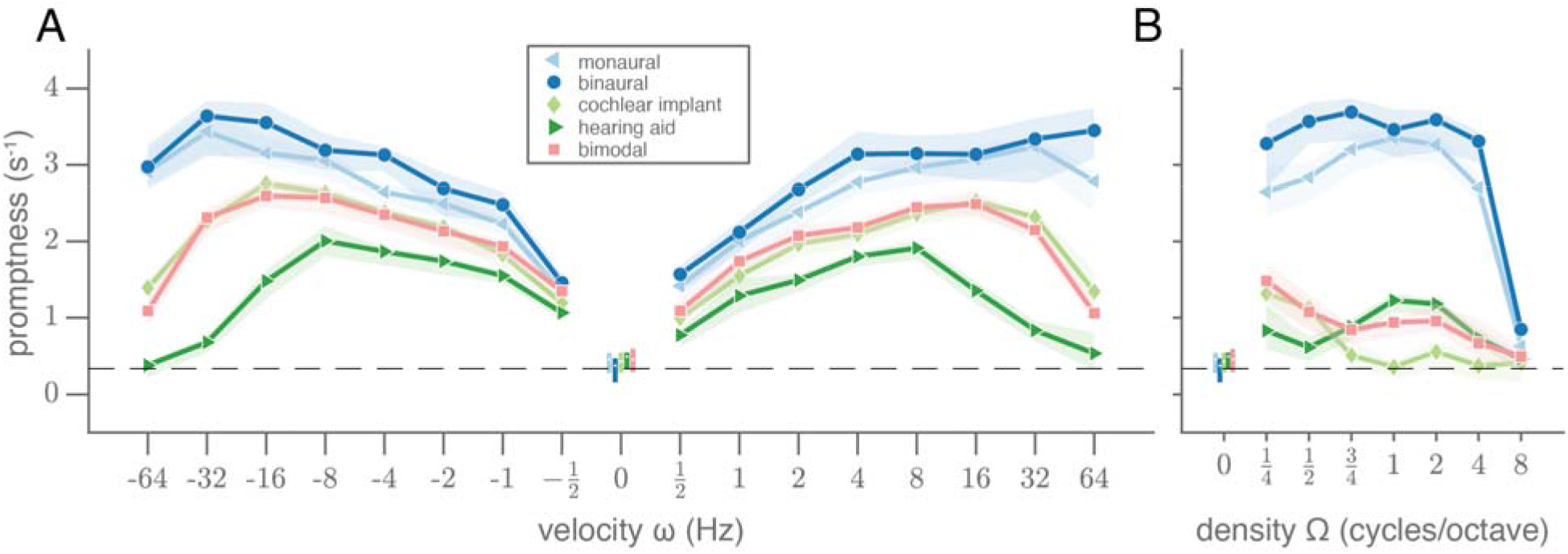
Pure temporal and spectral modulation transfer functions. A) Mean promptness (line and markers) and 95% confidence interval of the mean across subjects (patch) as a function of velocity (density = 0 cycles/octave) and B) as a function of density (velocity = 0 Hz), for each listening condition: normal-hearing monaural and binaural conditions, and the impaired-hearing cochlear-implant, hearing-aid and bimodal simulations. White dots and colored vertical lines around 0 indicate the mean promptness and its 95% confidence interval for the catch trials with unmodulated stimuli ([velocity, density] = [0 Hz, 0 cycles/octave]).

### Spectral-only modulations

For the static ripples (purely spectral modulations at velocity = 0 Hz), the promptness as a function of density resembled a low-pass characteristic, at least for the binaural and monaural normal-hearing conditions (Fig. 3B, dark blue circles and light blue triangles, respectively). For these conditions, detection is very poor for the highest density of 8 cycles/octave. This property presumably reflects the limits of resolvable power of the human auditory filters, leading to a poorer detection of spectral patterns with finer spectral detail.

Responses made for the cochlear-implant simulation (light green diamonds) resembled a band-pass filter characteristic with a cutoff around 0.75 cycles/octave and responses in the hearing-aid condition (dark green triangles) followed a band-pass characteristic with its highest promptness around 1-2 cycles/octave. Bimodal responses (pink squares) resembled the best values of the cochlear-implant and hearing-aid conditions. Overall, the impact of hearing-impairment simulation on reaction times was generally larger for the spectral modulations (Fig. 3B) than for the temporal modulations (Fig. 3A). This behavioral finding seems in line with the acoustic transformation effects of the vocoders on the sounds, that preserve temporal modulations to some extent (cf. Fig. 1, top row), but heavily perturb spectral modulations (cf. Fig. 1, side columns).

### Joint spectrotemporal modulation

Figs. 4A-E show the stMTFs for the two normal-hearing conditions, and for the three impaired-hearing simulations, for all joint spectrotemporal ripples, as mean promptness (averaged across listeners; see supplemental Figs. S1-6 for individual stMTFs) per ripple density (abscissa) and velocity (ordinate). Deep red colors correspond to high spectral-temporal sensitivity, dark blue colors to low sensitivity (low promptness values). The results for pure amplitude-modulated stimuli (cf. Fig. 3A) are at the bottom row of the stMTF matrix, at Ω=0 cycles/octave; the results for pure spectral modulations (cf. Fig. 3B) are found along the central column, at ω=0 Hz.

**Fig. 4.**
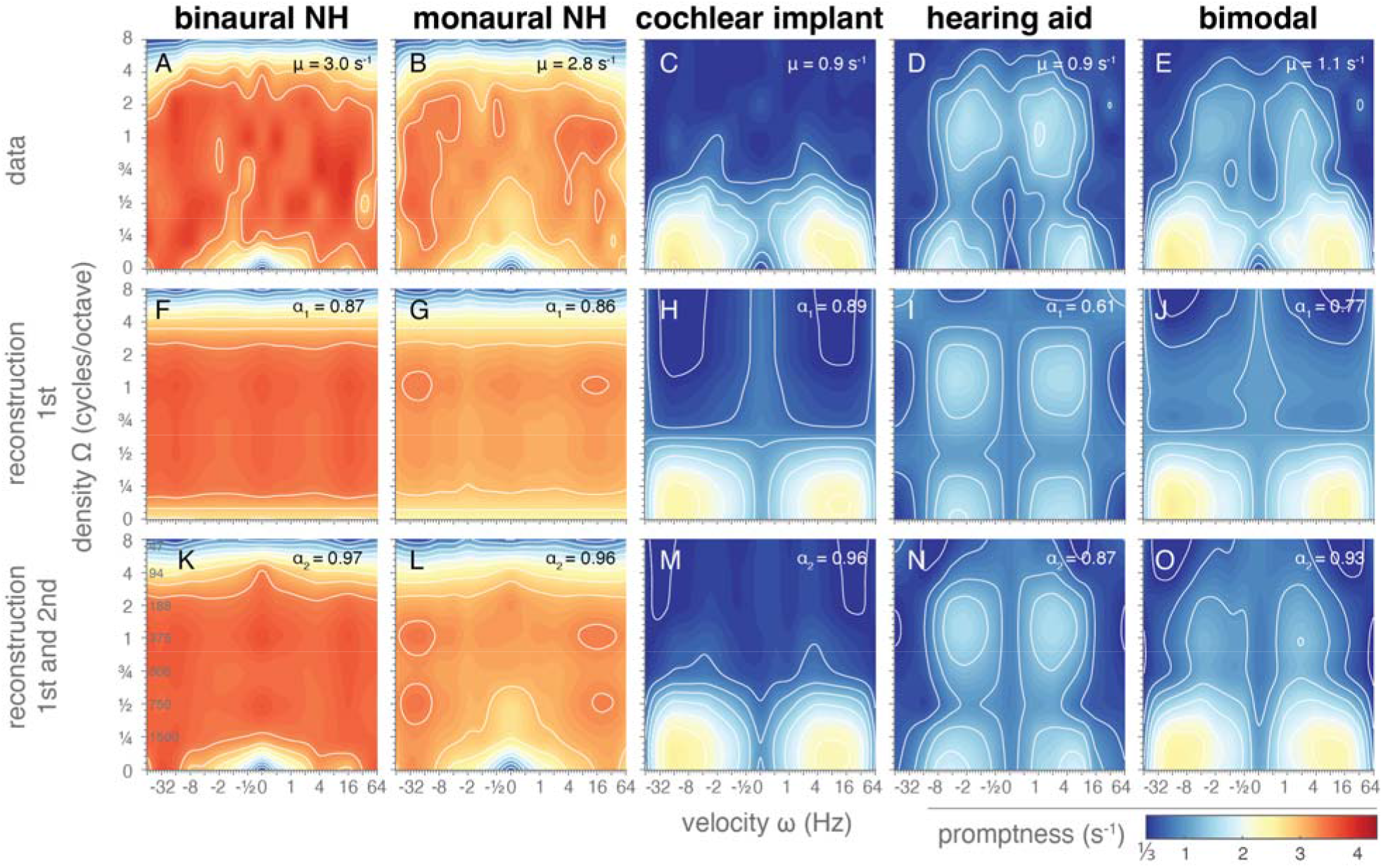
Spectrotemporal modulation transfer functions. Mean promptness as a function of velocity and density, representing spectrotemporal sensitivity in the normal hearing and simulated hearing-impaired conditions. F-J) Reconstructions of the stMTFs using only the first singular value after singular value decomposition (SVD; Eq. 4). K-O) Reconstructions of the stMTFs using the first and second SVD components (Eq. 6).

Ripple detection in the binaural normal-hearing condition (Fig. 4A) was faster (mean promptness = 3.0 s^-1^) than in the monaural condition (Fig. 4B; mean promptness: 2.8 s^-1^). Combined spectrotemporal modulation sensitivity for both listening conditions resembled a two-dimensional band-pass for both density and velocity, with fastest reaction times around velocities of ±8-16 Hz and densities around 0.75-1 cycles/octave (binaural maximum promptness = 4.0 s^-1^ at [ω, Ω]=[8 Hz, 0.75 cycles/octave]; monaural maximum promptness = 3.7 s^-1^ at [ω, Ω]=[-16 Hz, 1 cycles/octave]). The ripples were well detectable up to and including 4 cycles/octave.

The stMTFs for the impaired-hearing simulations (Fig. 4C-E) were distinctly slower when compared to normal hearing (mean promptness = 0.9, 0.9 and 1.1 s^-1^ for cochlear implant, hearing aid, and bimodal vocoder, respectively). Temporal modulation sensitivity again showed a band-pass filter characteristic with fastest detection rates around ±16 Hz. Cochlear-implant simulations (Fig. 4C) mainly reduced the detection of high spectral modulations, which is consistent with the modus operandi of a cochlear implant (and vocoders), whereby its band-pass filtering mechanism reduces spectral modulation sensitivity. The fastest responses (maximum promptness = 2.8 s^-1^) were elicited to -16 Hz amplitude-modulated sounds (Ω = 0 cycles/octave). Ripple detection with the cochlear-implant vocoder became impossible for densities exceeding 0.75 cycles/octave. Of all hearing conditions, listeners reacted slowest for the hearing-aid simulations (Fig. 4D; maximum promptness = 2.0 s^-1^). However, in contrast to the cochlear-implant condition, higher densities of up to 4 cycles/octave could still be detected if temporal modulation rates were not too fast (>16 Hz). A local dip in promptness exists for ripples with a density around 0.5 cycles/octave.

The bimodal simulation resembled a conjunction of the cochlear-implant and hearing-aid simulation results, seemingly exhibiting a ‘best of both worlds’ principle (Corneil et al., 2002) with responses almost as fast as for the best unimodal condition (Fig. 4E; maximum promptness = 2.6 s^-1^ at [ω, Ω]=[-16 Hz, 0 cycles/octave]). For high spectral modulation frequencies, the bimodal condition was comparable to the hearing-aid condition; for low spectral modulations it followed the cochlear-implant condition.

### Separability

We assessed the degree of separability of the stMTF into a pure temporal and spectral component through singular value decomposition (SVD) using the separability index α_1_ (Eq. 5) and the α_2_ index (Eq. 7) to determine the relative contribution of the first (Eq. 4) and of the first two components (Eq. 6). If the α_1_ index is close to 1, the MTF is considered to be separable (for individual-level separability indices and confidence intervals, see supplemental Fig. S7).

The central row of Fig. 4 shows the reconstructed stMTF_1_ for the various hearing conditions. The first-order reconstruction of the stMTF yielded purely orthogonal patterns in the matrix, resulting from the full-separability assumption. For both normal hearing conditions, the separability index α_1_ was high (0.87 and 0.86 for binaural and monaural listening conditions), which suggests that the variability in the normal-hearing stMTFs can be captured quite well with a first-order approximation (Figs. 4F,G) and that the matrix is highly separable (eq. 4). Notably missing in the reconstructions are the slow responses to the slow amplitude-modulated sounds (ω<1 Hz, Ω=0 cycles/octave; cf. Figs. 4A,B and Figs. 4F,G at the bottom of the images near the center). By adding the second singular value with its spectral and temporal components (bottom row of Fig. 4), the stMTF reconstructions improved considerably: α_2_=0.96 (monaural) and 0.97 (binaural). Now, the responses to the amplitude-modulated sounds seemed to be accounted for as well.

The separability index was best for the cochlear-implant simulation (Fig. 4H), which equaled or was better than for the normal-hearing conditions (α_1_=0.89). The first-order reconstructions were worse for the hearing-aid and bimodal conditions (α_1_= 0.61 and 0.77, respectively), suggesting a considerable inseparable spectrotemporal component to the responses of these two listening conditions. Incorporating the first two SVD components improved reconstructions (Figs. 4M-O): α_2_=0.96, 0.87, 0.93.

### Race models

To investigate to what extent monaural reaction times can predict binaural performance, we used the race model of statistical facilitation, which postulates independence between ears. As an example, Fig. 5A displays the cumulative reaction time probability of listener L6 for a stimulus with a 2 Hz, 0 cycles/octave modulation, for the monaural and binaural normal-hearing conditions, as well as for the promptness that would be reached based on the race model of statistical facilitation. For this ripple, binaural performance was faster than both the monaural condition and the race model. For the simulated listening conditions (Fig. 5B), the bimodal responses to this ripple were faster than the hearing-aid data and resembled the cochlear-implant data and the race model.

**Fig. 5.**
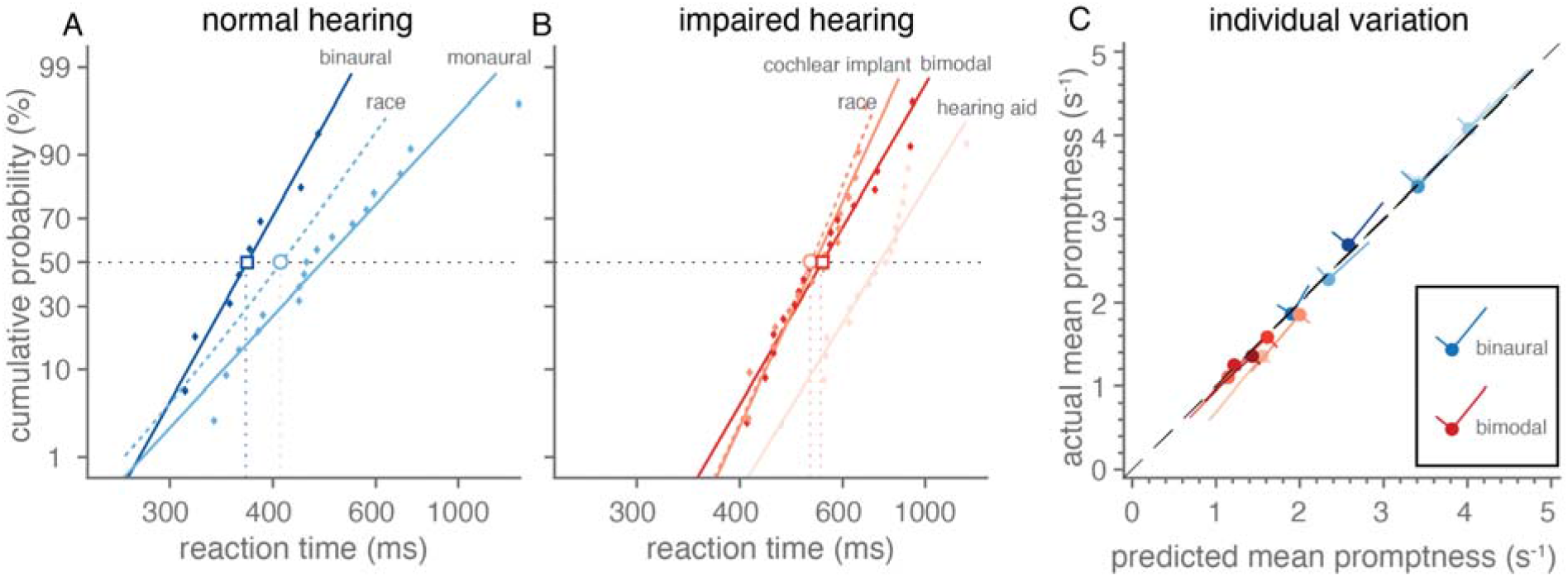
Binaural versus monaural detection of spectrotemporal modulations. A) Reaction time distributions of listener L6 plotted in the same format as in Figs. 2A-C for a stimulus with a 2 Hz, 0 cycles/octave modulation while listening monaurally (light blue) or binaurally (dark blue). The race model (determined from the monaural distribution according to Eqn. 8) is indicated by a light blue dashed line. Dots indicate individual reaction times; lines indicate the best linear fit through the data. The horizontal dotted line is drawn at a cumulative probability of 0.5 crossing the median reaction times. These medians for the bimodal data and the race model are indicated by an open square and circle, respectively. B) The same as in A, but now for the cochlear-implant, hearing-aid and bimodal listening conditions. Note that the binaural data is slightly faster than the race model, while bimodal data resembles both the cochlear-implant data and the race model. C) The median promptness as observed in the data is plotted as a function of the race-model prediction (closed circles and unbroken lines). Blueish and reddish colors indicate binaural and bimodal listening conditions, respectively. For each listener (indicated by different tint), the mean across ripple modulations is indicated by the closed circles, and standard deviation in the direction of the two axes with largest variability is indicated by lines. Note that the data fall closely on the unity-line (black dashed line), which holds both for the mean and the main axis with largest variability.

To quantify this for all ripples and listeners, we compared the median predictions from the benchmark race model to the median data (Fig. 5C). Overall, listeners were as fast as the race model prediction, both for binaural (Fig. 5C, blue circles on the diagonal) and bimodal hearing (Fig. 5C, reddish colors). These results show that binaural and bimodal performance seemed to follow statistical facilitation (Eqn. 8).

## Discussion

### Summary

This study used a speeded-response paradigm to determine the auditory stMTF in human listeners. The reaction times obtained appeared to be a valid and effective objective measure for ripple sensitivity, given its systematic relationship with the parameters that determine both temporal and spectral modulation rates. Sensitivity was highest for ripples with modulations around 16 Hz and 1 cycle/octave and decreased for higher and lower modulation rates. Using simulations of a cochlear-implant, hearing-aid and bimodal restorative hearing, spectrotemporal sensitivity worsened (reaction times increased) compared to normal hearing, as expected from the impaired signal processing of the simulations. Although the separability of the stMTF into a spectral and temporal component was high for both the normal-hearing and for the cochlear-implant simulated data, the inseparable second-order spectrotemporal component was still substantial, with a value between 7-10%. For the bimodal and especially the hearing-aid conditions, inseparability was larger by about 16 and 28%. For all ripples, binaural and bimodal reaction-time performance was comparable to the prediction of the benchmark race model of statistical facilitation, suggesting independent detection of two monaural signals, rather than true binaural integration.

### Normal Hearing

Constructing the stMTF based on reaction times is a fairly new approach that has been introduced so far only in monkey research (Massoudi et al. 2013, 2014). Still, the observed stMTFs (Fig. 3 and Fig. 4A-E) correspond well with the results of other studies in humans, which have measured modulation detection thresholds. Chi et al. (1999) and Zheng et al. (2017) measured the full stMTF for normal-hearing listeners using an adaptive modulation-detection threshold paradigm and found band-pass functions for both the spectral and temporal dimensions. They found best ripple detection-thresholds at spectral modulations below or at 1 cycle/octave, and temporal modulations around 4-16 Hz. Despite these small quantitative differences between studies, the general patterns were similar, and in line with our results (Fig. 6): when comparing our promptness data (Fig. 6C,D) with the modulation thresholds collected in the earlier studies (Chi et al. 1999; Zheng et al. 2017 in Fig. 6A,B), the stMTFs resemble each other, at least qualitatively. These convergent findings suggest that reaction times are indeed a valid objective measure to determine the spectrotemporal sensitivity of (naïve) listeners.

**Fig. 6.**
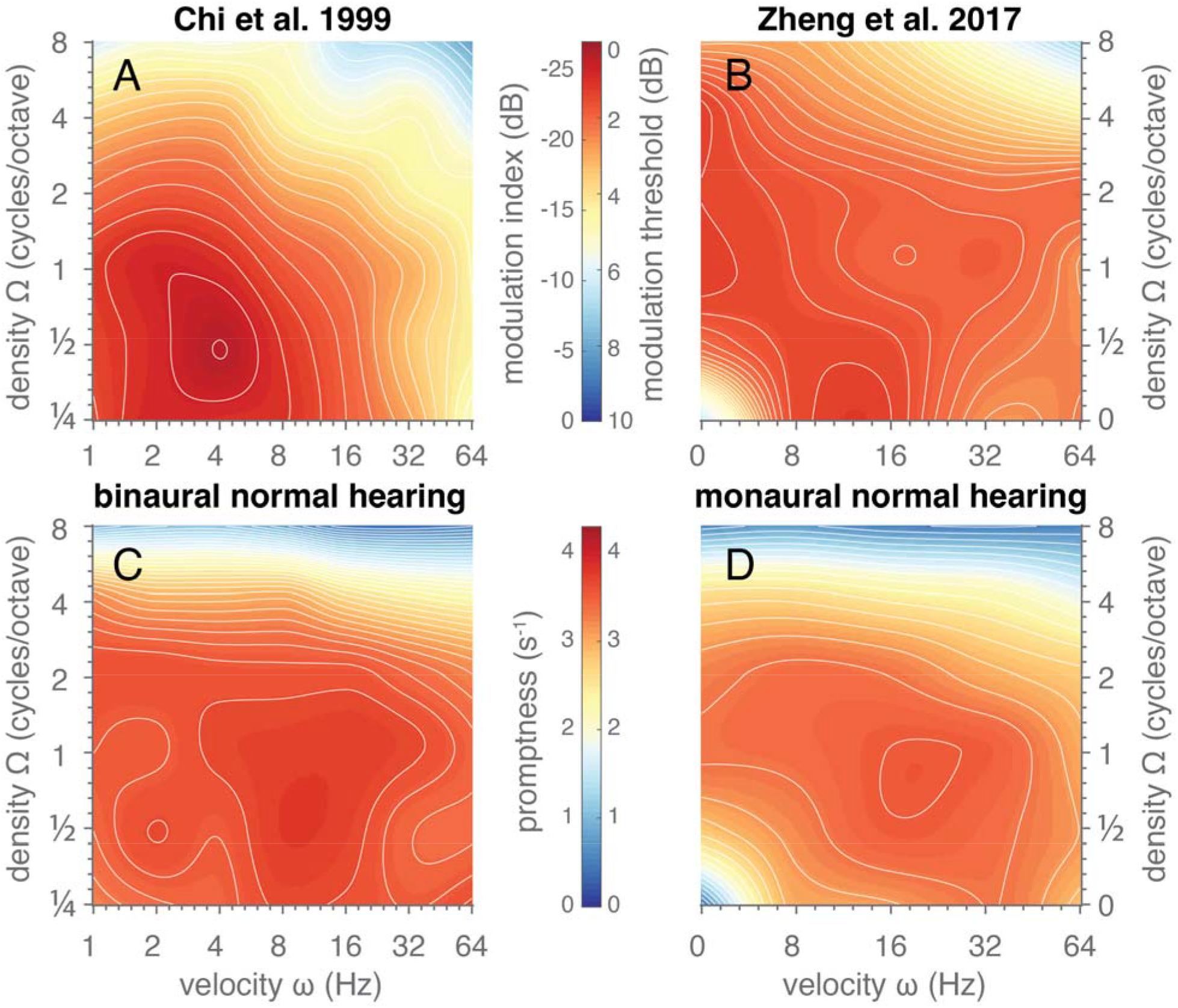
Comparison of spectrotemporal sensitivity. Images depict stMTFs obtained using **A**,**B)** a threshold-searching paradigm from the studies of **A)** Chi et al. 1999 and B) Zheng et al. 2017 or **C**,**D)** a reaction-time task from this study in the **C)** binaural and **D)** monaural listening conditions. Data from previous studies was obtained from Figs. 3 of the respective papers. Image format is the same as in Fig. 4. Data (modulation index) in **A)** was log-transformed (20log_10_) to match the data (modulation threshold in dB) in **B)**. Data from this study in C and D was replotted from Fig. 4, matching the velocities and densities used in the previous studies (A vs C, B vs D). Note that the colour scales in A,B) are reversed in order, as best responses correspond to high promptness values, but low modulation indices or thresholds.

Chi et al. (1999) proposed a computational model to explain their data, in which the spectrotemporal modulation sensitivity is based on cortical responses to the ripple’s spectrogram. The modulation transfer functions generated by their model closely resembled their data, and thus will resemble our data as well. They concluded that ‘the upper limits of the spectral and temporal modulation rates are related through the effective bandwidths of the cochlear filters’ (Chi et al. 1999).

Narne et al. (2016, 2018, 2020) have studied spectral resolution by means of a spectral-ripple or a moving-ripple test. They found thresholds around 5 to 6 cycles/octave for normal-hearing listeners in optimal conditions. Again, this seems to be in line with the strong drops in sensitivity observed in our data for densities at 8 cycles/oct in the non-processed conditions. This also suggests that we might have missed a more gradual decline of promptness, as we did not study any densities between 4 and 8 kHz.

### Impaired-hearing simulations

To evaluate our reaction-time test under more challenging listening situations, we manipulated the ripple stimuli using hearing-aid and cochlear-implant simulations. Both simulations made it substantially harder to detect ripple modulations and even impossible for certain parameters, eliciting much longer reaction times compared to monaural normal hearing for all ripples. Bimodal hearing exhibited a ‘best-of-both-worlds’ effect, following the fastest unimodal condition, which was the cochlear-implant condition for spectral modulations below 0.75 cycles/octave, and the hearing aid for higher spectral modulations (Fig. 4). An improvement in spectral ripple discrimination for bimodal hearing over the cochlear-implant alone has also been found in users with combined electro-acoustic stimulation in the same ear (Golub et al. 2012).

Other studies have shown a 5-10 dB reduction in the temporal-modulation detection threshold for cochlear-implant users compared to normal hearing (Bacon and Viemeister 1985; Golub et al. 2012; Won et al. 2011), whereas we found a decrease in promptness of 0.4 ± 0.2 s^-1^ for the well-detectable rates below 16 Hz. In these studies, hearing-impaired listeners performed in between normal hearing and cochlear-implant users. Our hearing-aid simulation, however, showed longer reaction times for temporal modulations (at 0 cycles/octave) compared to the cochlear implant condition. It should be emphasized that our hearing-aid condition was based on a worst-case-scenario, simulating very little residual hearing, whereas hearing thresholds of the hearing-impaired listeners in the study of Bacon and Viemeister (1985) still reached up to 10-20 dB HL at 1 kHz. Their study also showed a link between degraded temporal sensitivity and reduced listening bandwidth.

Impaired spectral modulation sensitivity with a cochlear implant is a likely result of its band-pass filtering mechanism that limits the spectral information to a set number of spectral bands. Henry et al. (2005), Berenstein et al. (2008) and Narne et al. (2020) all found lower spectral ripple modulation thresholds for cochlear-implant users compared to normal-hearing listeners, roughly corresponding to the increased reaction times in our study. Spectral modulation thresholds in hearing-impaired listeners have been reported to be 5-10 dB worse than for normal hearing (Davies-Venn, Nelson, and Souza 2015; Summers and Leek 1994), which may agree with the longer reaction times of our hearing-aid simulation compared to normal hearing. A few studies investigated combined spectrotemporal modulation detection thresholds in hearing-impaired listeners, which were often worse compared to normal hearing listeners, especially for low temporal modulation rates (Bernstein et al. 2013; Mehraei et al. 2014; Zheng et al. 2017; Zhou et al. 2020).

### Race model

To get insights in the mechanism of combining input at both ears, we used race models to test whether monaural responses could predict binaural performance. For normal-hearing conditions, faster reaction times were elicited when stimuli were presented binaurally compared to monaural presentation (Fig. 4A,B). Binaural responses seemed as fast as the race model of statistical facilitation (Fig. 5C). This suggests that ripple detection was determined by a parallel race between the two ears, rather than from neural integration.

Bimodal responses were also as fast as the race model (Fig. 5C), suggesting that there is also no benefit of integration for the poorest listening conditions. This is interesting if we compare this finding to audiovisual gaze-orienting experiments that aim to study neural integration of visual and auditory signals. Strongest benefits of multisensory interactions (i.e., increased speed, accuracy, and precision of responses) are obtained for stimuli that overlap both in space and time, and thus provide multisensory evidence for a single object. Moreover, these interactions are strongest when the uni-sensory evoked responses are variable and slow (i.e., away from ceiling performance). This phenomenon is known as the ‘principle of inverse effectiveness’ (e.g. Corneil et al. 2002; van de Rijt et al. 2019; Stein and Meredith 1993; Van Wanrooij et al. 2009). We here propose that beneficial effects of bimodal (cochlear-implant-hearing-aid) integration will depend on whether the auditory system has sufficient evidence that left vs. right acoustic inputs arose from the same auditory object, rather than from unrelated sounds. The strongest bimodal benefits (i.e., enhanced sensitivity) will thus be found: (i) when spectral ranges of cochlear implant and hearing-aid overlap sufficiently (for within-spectral comparisons), and (ii) when monaural reaction-time distributions have sufficient variability and overlap considerably. Since neither of these two requirements seem to be fulfilled, it may be unsurprising that bimodal listening does not exceed race-model performance and does not benefit from neural integration. Instead, the benefit of bimodal listening seems to consist purely of predominantly perceiving low-density temporal modulations with the cochlear-implant ear and low-velocity spectral modulations with the hearing-aid ear (Figs. 4C-E).

In contrast to our results, several studies have shown reaction times to stimuli that exceeded the predictions based on statistical facilitation. However, these studies typically involved responses to multisensory stimuli, or to the dichotic presentation of two spectrally distinct sounds (Gielen et al. 1983; Miller 1982; Schröter et al. 2007; Townsend and Nozawa 1995). Like the findings of Schröter et al. (2007) for auditory stimuli that fused into a single percept, we did not obtain faster responses than expected from statistical facilitation in the bimodal conditions. The benefits of an integrative process must be found outside the parameters presented in this study, and likely include the ability to localize sounds and enhancement of speech perception in noisy environments.

### Separability

Measuring the stMTF is typically a time-consuming process, for which it would be valuable to know whether the two-dimensional function is simply the product of a temporal and spectral component. A large degree of separability of spectrotemporal sensitivity has been found for normal-hearing listeners and cochlear-implant users (Chi et al. 1999; Zheng et al. 2017). Our data support this notion of large separability (but not ‘full’ separability) for the normal-hearing and cochlear-implant listening conditions. Nevertheless, the contribution of the second singular value was still substantial, close to 10% in the normal-hearing condition. For the hearing-aid and bimodal listening conditions, the contribution could be as large as 20%, implying a large degree of inseparability. Studies by Bernstein et al. (2013) and Zheng et al. (2017) found that spectrotemporal cues may enhance speech intelligibility and modulation sensitivity in CI users over spectral or temporal modulations alone. These findings hint at some form of spectrotemporal integration, but interestingly this may not be captured by the relatively large separability of the stMTF (Zheng et al. 2017).

As can be appreciated from the observable differences between the single and two-component reconstructions (Fig. 4), the second component adds a diagonal interaction component to the two-dimensional MTF. Most of this interaction yields changes to the MTF at 0- or low-density modulations: reactions become slower for low-velocity modulations, and faster for higher velocities. We conclude that a full assessment of spectrotemporal performance requires testing the complete or large spectrotemporal space of all ripples, by including the second SVD component (Fig. 4).

### A novel hearing test?

We believe that a reaction-time paradigm could provide an alternative novel hearing test. This test potentially provides a description of acoustic sensitivity that is potentially closer to speech perception accuracy than pure-tone audiometry. Here, we demonstrated the convergent validity of the test by showing that the stMTF obtained from reaction times resemble the stMTFs obtained from standard detection thresholds (Fig. 6). Earlier studies have shown that reaction times (chronometric function) and detection thresholds (psychometric function) are tightly coupled for certain experimental paradigms (Palmer, Alexander C. Huk, and Shadlen 2005). By using sequential-sampling or drift-diffusion models (Palmer, Alexander C Huk, and Shadlen 2005; Rach, Diederich, and Colonius 2011), the two measures could be integrated into a single framework.. A study on audiovisual integration shows that the two measures provide complementary, but not the same information (Rach et al. 2011) and a study on pure-tone detection also found the measures to be not interchangeable (Abel, Rajan, and Giguere 2009). How strong the coupling is for the modulation-onset detection test is of yet unknown.

The test is reliable across listeners, yielding highly similar stMTFs (except for an idiosyncratic reaction time offset, supplemental Figs. S1-6) and separability indices with narrow confidence intervals (supplemental Fig. S7). An advantage over a modulation-detection threshold paradigm is that stimuli can be presented at supra-threshold levels, allowing for a relatively easy task, potentially suitable for clinical assessments, and for studies with children, or with experimental animals. This is supported by the fact that we did not observe procedural learning effects during the experiments.

Nevertheless, usability and construct validity of a reaction-time hearing test should be explored further. There is reason to believe that stimulus parameters other than velocity and density may influence the stMTF. Modulation depth, sound level, and spectral profile each may affect the stMTF by enhancing or suppressing modulation sensitivity. For example, a full modulation depth of 100% may produce a richer characterization of the modulation space especially in the impaired conditions and reduce the number of conditions in which the stimuli are undetectable. Similarly, by having a carrier with a speech-shaped spectrum or a pink spectrum, detection of modulation onset might correlate better with detection of natural sounds, such as speech (phonemes or words) or music. In fact, one can think of developing a framework that relates the sensitivity to the spectrotemporal modulations of the moving ripples to the recognition and pitch accuracy of speech materials such as phonemes and words (Elliott and Theunissen 2009). While beyond the scope of this paper, note that theoretically these dynamic signals can be decomposed into their constituent ripples. In principle, there could be a direct relationship, although confounded by noise in the responses and by cognitive, non-acoustic aspects (such as predictability of a word in a sentence).

Numerous studies have shown that non-acoustic aspects, such as attention and reward expectation, could affect the spectrotemporal receptive fields of cells in primary auditory cortex (Atiani et al., 2009; Bakin & Weinberger, 1996; David et al., 2012; Fritz et al., 2003; Fritz et al., 2005a,b; Fritz et al., 2007; Jaramillo & Zador, 2011; Ji et al., 2001; Kilgard et al., 2001, 2002; Kilgard & Merzenich, 2002; Lee & Middlebrooks, 2011; Ohl & Scheich, 1997, 2005; Suga et al., 2002; Suga & Ma, 2003). It would not be farfetched to believe that these changes would also be reflected in the overt behavior. In our experiments, we chose to randomize the order of listening conditions and stimulus parameters, making it impossible for the listener to predict what is coming up. This procedure allows for a direct, acute assessment of acoustic spectrotemporal sensitivity. The downside, of course, is that any adaptive change in the stMTF will not be captured by this test. Conducting the same task following a blocked design, might produce different filters. Similarly, the stMTFs observed for the cochlear-implant and hearing-aid simulations might differ from those that would be obtained from actual hearing-impaired listeners. Due to long-term experience, they could have acclimatized or adapted to their hearing impairment.

## Conclusion

Reaction times are a valid objective measure for ripple sensitivity. The joint spectrotemporal transfer function closely resembled data from earlier studies that used modulation detection thresholds. Responses to spectrotemporal modulated ripples could be reconstructed by using the first two components of singular value decomposition, suggesting significant spectrotemporal inseparability, especially for the hearing-aid and bimodal listening conditions. We further found that binaural and simulated bimodal reaction times could be predicted from statistical facilitation induced by a race of independent monaural inputs.

## Acknowledgements

We thank Ruurd Lof, Stijn Martens, and Gu□nter Windau for their technical support and Daisy Louvet for her assistance in data collection.

## Data availability statement

Data are available from the Donders Institute for Brain, Cognition and Behaviour repository at https://data.donders.ru.nl and can be found via the following persistent identifier: https://doi.org/10.34973/87nw-zb11.

## References

Abel, S. M., D. K. Rajan, and C. Giguere. 2009. “Stimulus Parameters in Detection and Reaction Time.” http://Dx.Doi.Org/10.3109/01050399009070777 19(4):229–35.

Ausili, S. A., B. Backus, M. J. H. Agterberg, A. J. van Opstal, and M. M. van Wanrooij. 2019. “Sound Localization in Real-Time Vocoded Cochlear-Implant Simulations With Normal-Hearing Listeners.” Trends in Hearing 23.

Bacon, S. P., and N. F. Viemeister. 1985. “Temporal Modulation Transfer Functions in Normal-Hearing and Hearing-Impaired Listeners.” Audiology□: Official Organ of the International Society of Audiology 24(2):117–34.

Baer, Thomas, and Brian C. J. Moore. 1994. “Effects of Spectral Smearing on the Intelligibility of Sentences in the Presence of Interfering Speech.” The Journal of the Acoustical Society of America 95(4):2277.

Berenstein, Carlo K., Lucas H. M. Mens, Jef J. S. Mulder, and Filiep J. Vanpoucke. 2008. “Current Steering and Current Focusing in Cochlear Implants: Comparison of Monopolar, Tripolar, and Virtual Channel Electrode Configurations.” Ear and Hearing 29(2):250–60.

Bernstein, Joshua G. W., Golbarg Mehraei, Shihab Shamma, Frederick J. Gallun, Sarah M. Theodoroff, and Marjorie R. Leek. 2013. “Spectrotemporal Modulation Sensitivity as a Predictor of Speech Intelligibility for Hearing-Impaired Listeners.” Journal of the American Academy of Audiology 24(4):293–306.

Bernstein, Leslie R., and David M. Green. 1998. “Detection of Simple and Complex Changes of Spectral Shape.” The Journal of the Acoustical Society of America 82(5):1587.

Carpenter, R. H. S., B. A. J. Reddi, and A. J. Anderson. 2009. “A Simple Two-Stage Model Predicts Response Time Distributions.” The Journal of Physiology 587(16):4051–62.

Carpenter, R. H., and M. L. Williams. 1995. “Neural Computation of Log Likelihood in Control of Saccadic Eye Movements.” Nature 377(6544):59–62.

Chi, Taishih, Yujie Gao, Matthew C. Guyton, Powen Ru, and Shihab Shamma. 1999. “Spectro-Temporal Modulation Transfer Functions and Speech Intelligibility.” The Journal of the Acoustical Society of America 106(5):2719–32.

Corneil, B. D., M. Van Wanrooij, D. P. Munoz, and a J. Van Opstal. 2002. “Auditory-Visual Interactions Subserving Goal-Directed Saccades in a Complex Scene.” Journal of Neurophysiology 88(1):438–54.

Davies-Venn, Evelyn, Peggy Nelson, and Pamela Souza. 2015. “Comparing Auditory Filter Bandwidths, Spectral Ripple Modulation Detection, Spectral Ripple Discrimination, and Speech Recognition: Normal and Impaired Hearinga).” The Journal of the Acoustical Society of America 138(1):492.

Elliott, Taffeta M., and Frédéric E. Theunissen. 2009. “The Modulation Transfer Function for Speech Intelligibility” edited by K. J. Friston. PLoS Computational Biology 5(3):e1000302.

Gielen, Stan C. A. M., Richard A. Schmidt, and Pieter J. M. Van Den Heuvel. 1983. “On the Nature of Intersensory Facilitation of Reaction Time.” Perception & Psychophysics 1983 34:2 34(2):161–68.

Glasberg, Brian R., and Brian C. J. Moore. 1986. “Auditory Filter Shapes in Subjects with Unilateral and Bilateral Cochlear Impairments.” The Journal of the Acoustical Society of America 79(4):1020.

Golub, Justin S., Jong Ho Won, Ward R. Drennan, Tina D. Worman, and Jay T. Rubinstein. 2012. “Spectral and Temporal Measures in Hybrid Cochlear Implant Users: On the Mechanism of Electroacoustic Hearing Benefits.” Otology and Neurotology 33(2):147–53.

Henry, Belinda A., and Christopher W. Turner. 2003. “The Resolution of Complex Spectral Patterns by Cochlear Implant and Normal-Hearing Listeners.” The Journal of the Acoustical Society of America 113(5):2861.

Henry, Belinda A., Christopher W. Turner, and Amy Behrens. 2005. “Spectral Peak Resolution and Speech Recognition in Quiet: Normal Hearing, Hearing Impaired, and Cochlear Implant Listeners.” The Journal of the Acoustical Society of America 118(2):1111–21.

Hershenson, Maurice. 1962. “Reaction Time as a Measure of Intersensory Facilitation.” Journal of Experimental Psychology 63(3):289–93.

Liberman, Alvin M. (Alvin Meyer). 1996. “SpeechL: A Special Code.” 458.

Litvak, Leonid M., Anthony J. Spahr, Aniket A. Saoji, and Gene Y. Fridman. 2007. “Relationship between Perception of Spectral Ripple and Speech Recognition in Cochlear Implant and Vocoder Listeners.” The Journal of the Acoustical Society of America 122(2):982.

Loizou, Philipos C., Michael Dorman, and Zhemin Tu. 1999. “On the Number of Channels Needed to Understand Speech.” The Journal of the Acoustical Society of America 106(4):2097.

Massoudi, Roohollah, Marc M. Van Wanrooij, Huib Versnel, and A. John Van Opstal. 2015. “Spectrotemporal Response Properties of Core Auditory Cortex Neurons in Awake Monkey” edited by M. S. Malmierca. PLOS ONE 10(2):e0116118.

Massoudi, Roohollah, Marc M. Van Wanrooij, Sigrid M. C. I. Van Wetter, Huib Versnel, and A. John Van Opstal. 2013. “Stable Bottom-up Processing during Dynamic Top-down Modulations in Monkey Auditory Cortex.” European Journal of Neuroscience (February):n/a-n/a.

Massoudi, Roohollah, Marc M. Van Wanrooij, Sigrid M. C. I. Van Wetter, Huib Versnel, and A. John Van Opstal. 2014. “Task-Related Preparatory Modulations Multiply with Acoustic Processing in Monkey Auditory Cortex.” European Journal of Neuroscience n/a-n/a.

McDermott, Josh H., and Eero P. Simoncelli. 2011. “Sound Texture Perception via Statistics of the Auditory Periphery: Evidence from Sound Synthesis.” Neuron 71(5):926–40.

Mehraei, Golbarg, Frederick J. Gallun, Marjorie R. Leek, and Joshua G. W. Bernstein. 2014. “Spectrotemporal Modulation Sensitivity for Hearing-Impaired Listeners: Dependence on Carrier Center Frequency and the Relationship to Speech Intelligibility.” The Journal of the Acoustical Society of America 136(1):301.

Miller, J. 1982. “Divided Attention: Evidence for Coactivation with Redundant Signals.” Cogn Psychol. 14(2):247–79.

Moore, B. C., and B. R. Glasberg. 2001. “Temporal Modulation Transfer Functions Obtained Using Sinusoidal Carriers with Normally Hearing and Hearing-Impaired Listeners.” The Journal of the Acoustical Society of America 110(2):1067–73.

Narne, Vijaya Kumar, Saransh Jain, Chitkala Sharma, Thomas Baer, and Brian C. J. Moore. 2020. “Narrow-Band Ripple Glide Direction Discrimination and Its Relationship to Frequency Selectivity Estimated Using Psychophysical Tuning Curves.” Hearing Research 389:107910.

Narne, Vijaya Kumar, Prashanth Prabhu, Bram Van Dun, and Brian C. J. Moore. 2018. “Ripple Glide Direction Discrimination and Its Relationship to Frequency Selectivity Estimated Using Notched Noise.” Acta Acustica United with Acustica 104(6):1063–74.

Narne, Vijaya Kumar, Mridula Sharma, Bram Van Dun, Shalini Bansal, Latika Prabhu, and Brian C. J. Moore. 2016. “Effects of Spectral Smearing on Performance of the Spectral Ripple and Spectro-Temporal Ripple Tests.” The Journal of the Acoustical Society of America 140(6):4298.

Palmer, John, Alexander C Huk, and Michael N. Shadlen. 2005. “And Accuracy of a Perceptual Decision.” 376–404.

Palmer, John, Alexander C. Huk, and Michael N. Shadlen. 2005. “The Effect of Stimulus Strength on the Speed and Accuracy of a Perceptual Decision.” Journal of Vision 5(5):1–1.

Raab, D. H. 1962. “Statistical Facilitation of Simple Reaction Times.” Transactions of the New York Academy of Sciences 24(5 Series II):574–90.

Rach, Stefan, Adele Diederich, and Hans Colonius. 2011. “On Quantifying Multisensory Interaction Effects in Reaction Time and Detection Rate.” Psychological Research 75(2):77–94.

Ratcliff, Roger, and Hans P. A. Van Dongen. 2011. “Diffusion Model for One-Choice Reaction-Time Tasks and the Cognitive Effects of Sleep Deprivation.” Proceedings of the National Academy of Sciences of the United States of America 108(27):11285–90.

van de Rijt, Luuk P. H., Anja Roye, Emmanuel A. M. Mylanus, A. John van Opstal, and Marc M. van Wanrooij. 2019. “The Principle of Inverse Effectiveness in Audiovisual Speech Perception.” Frontiers in Human Neuroscience 13:335.

Schröter, Hannes, Rolf Ulrich, and Jeff Miller. 2007. “Effects of Redundant Auditory Stimuli on Reaction Time.” Psychonomic Bulletin & Review 14(1):39–44.

Shannon, R. V, F. G. Zeng, V. Kamath, J. Wygonski, and M. Ekelid. 1995. “Speech Recognition with Primarily Temporal Cues.” Science (New York, N.Y.) 270(5234):303–4.

Sharma, Snandan, Lucas H. M. Mens, Ad F. M. Snik, A. John van Opstal, and Marc M. van Wanrooij. 2019. “An Individual With Hearing Preservation and Bimodal Hearing Using a Cochlear Implant and Hearing Aids Has Perturbed Sound Localization but Preserved Speech Perception.” Frontiers in Neurology 10:637.

Sharma, Snandan, Waldo Nogueira, A. John van Opstal, Josef Chalupper, Lucas H. M. Mens, and Marc M. van Wanrooij. 2021. “Amount of Frequency Compression in Bimodal Cochlear Implant Users Is a Poor Predictor for Audibility and Spatial Hearing.” Journal of Speech, Language, and Hearing Research 1–14.

Stein, B. E., and M. A. Meredith. 1993. The Merging of the Sense. Cambridge, MA, US: The MIT Press.

Summers, V., and M. R. Leek. 1994. “The Internal Representation of Spectral Contrast in Hearing-Impaired Listeners.” The Journal of the Acoustical Society of America 95(6):3518–28.

Supin, Alexander Ya, Vladimir V. Popov, Olga N. Milekhina, and Mikhail B. Tarakanov. 1994. “Frequency Resolving Power Measured by Rippled Noise.” Hearing Research 78(1):31–40.

Townsend, James T., and Georgie Nozawa. 1995. “Spatio-Temporal Properties of Elementary Perception: An Investigation of Parallel, Serial, and Coactive Theories.” Journal of Mathematical Psychology 39(4):321–59.

Versnel, Huib, Marcel P. Zwiers, and A. John van Opstal. 2009. “Spectrotemporal Response Properties of Inferior Colliculus Neurons in Alert Monkey.” The Journal of Neuroscience□: The Official Journal of the Society for Neuroscience 29(31):9725–39.

Veugen, Lidwien C. E., Josef Chalupper, Ad F. M. Snik, A. John van Opstal, and Lucas H. M. Mens. 2016. “Frequency-Dependent Loudness Balancing in Bimodal Cochlear Implant Users.” Acta Oto-Laryngologica 136(8):775–81.

Veugen, Lidwien C.E., Maartje M. E. Hendrikse, Marc M. van Wanrooij, Martijn J. H. Agterberg, Josef Chalupper, Lucas H. M. Mens, Ad F. M. Snik, and A. John van Opstal. 2016. “Horizontal Sound Localization in Cochlear Implant Users with a Contralateral Hearing Aid.” Hearing Research 336:72–82.

Viemeister, Neal F. 1998. “Temporal Modulation Transfer Functions Based upon Modulation Thresholds.” The Journal of the Acoustical Society of America 66(5):1364.

Van Wanrooij, Marc M., Andrew H. Bell, Douglas P. Munoz, and A. John Van Opstal. 2009. “The Effect of Spatial-Temporal Audiovisual Disparities on Saccades in a Complex Scene.” Experimental Brain Research 198(2–3):425–37.

Won, Jong Ho, Ward R. Drennan, Kaibao Nie, Elyse M. Jameyson, and Jay T. Rubinstein. 2011. “Acoustic Temporal Modulation Detection and Speech Perception in Cochlear Implant Listenersa).” The Journal of the Acoustical Society of America 130(1):376.

Zheng, Yi, Monty Escabí, and Ruth Y. Litovsky. 2017. “Spectro-Temporal Cues Enhance Modulation Sensitivity in Cochlear Implant Users.” Hearing Research 351:45–54.

Zhou, Ning, Susannah Dixon, Zhen Zhu, Lixue Dong, and Marti Weiner. 2020. “Spectrotemporal Modulation Sensitivity in Cochlear-Implant and Normal-Hearing Listeners: Is the Performance Driven by Temporal or Spectral Modulation Sensitivity?” Trends in Hearing 24:1–11.

