## supplemental Fig. for "Reaction time sensitivity to spectrotemporal modulations of sound"

Supplemental Material

### Individual spectrotemporal modulation transfer functions

Figs. S1-6 show the spectrotemporal modulation transfer functions for the two normal-hearing conditions, and for the three impaired-hearing simulations, for all joint spectrotemporal ripples, as mean promptness for all listeners individually. These figures correspond to Fig. 4 in the main document.

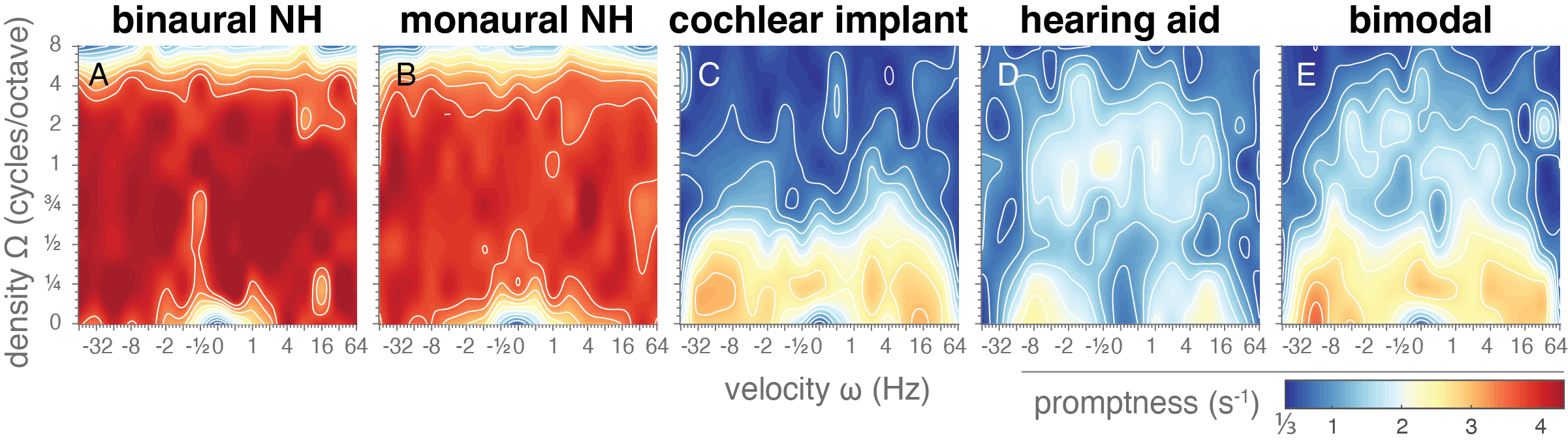

**Fig. S1. Spectrotemporal modulation transfer functions of listener L1.** Mean promptness as a function of velocity and density, representing spectrotemporal sensitivity in the normal hearing and simulated hearing-impaired conditions.

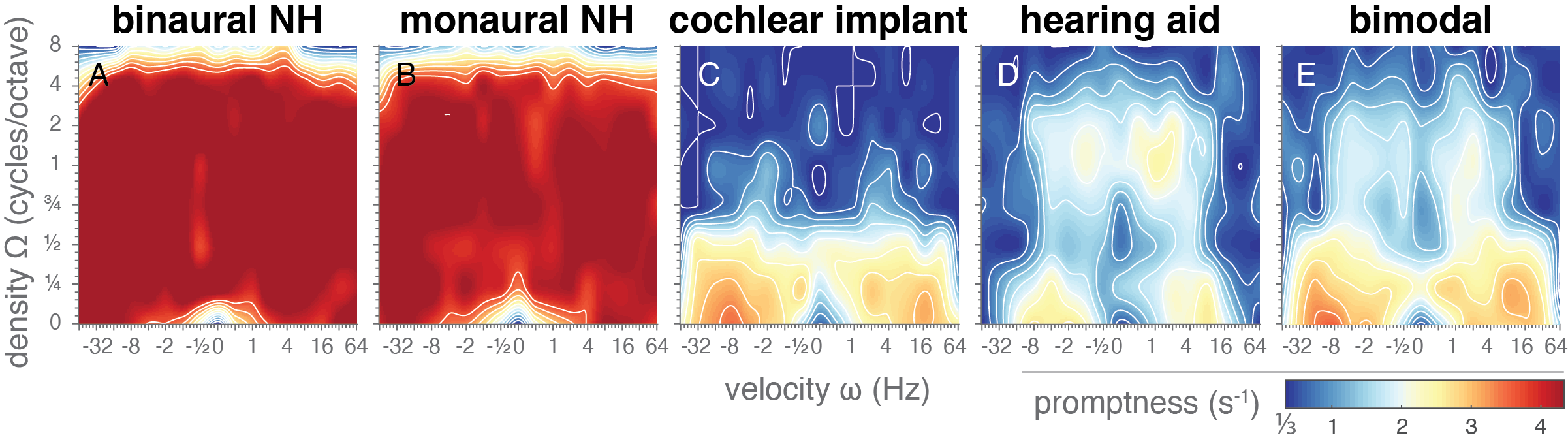

**Fig. S2. Spectrotemporal modulation transfer functions of listener L2.** Mean promptness as a function of velocity and density, representing spectrotemporal sensitivity in the normal hearing and simulated hearing-impaired conditions.

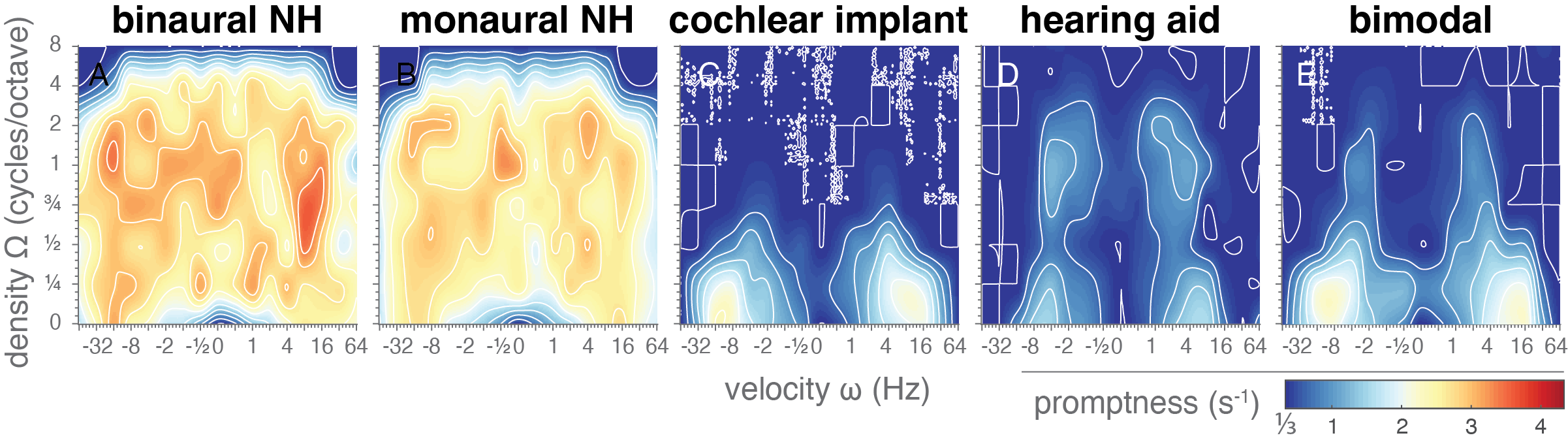

**Fig. S3. Spectrotemporal modulation transfer functions of listener L3.** Mean promptness as a function of velocity and density, representing spectrotemporal sensitivity in the normal hearing and simulated hearing-impaired conditions.

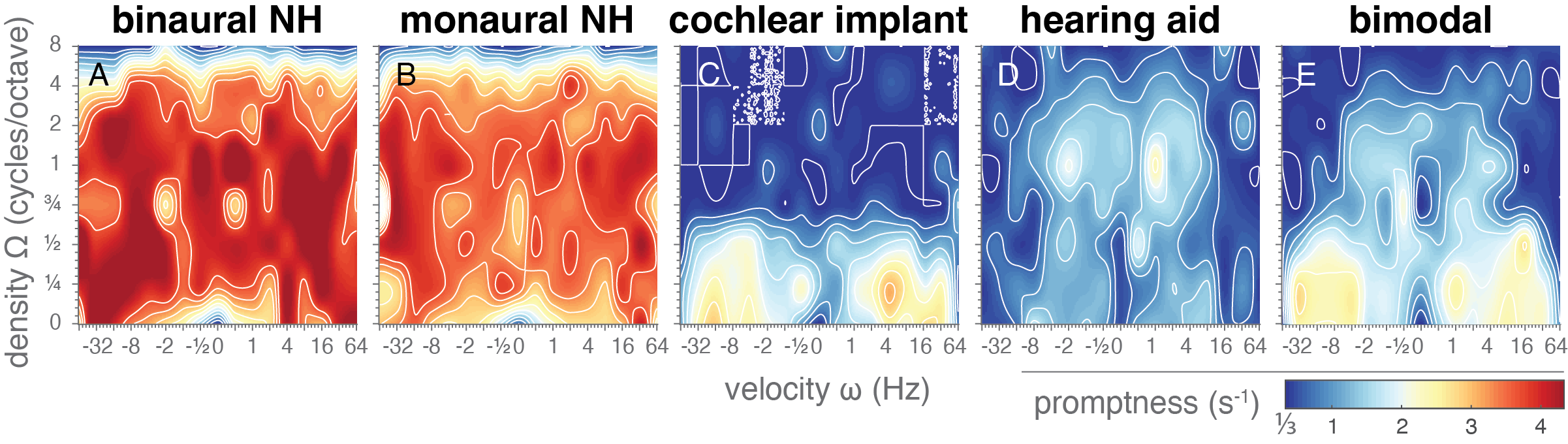

**Fig. S4. Spectrotemporal modulation transfer functions of listener L4.** Mean promptness as a function of velocity and density, representing spectrotemporal sensitivity in the normal hearing and simulated hearing-impaired conditions.

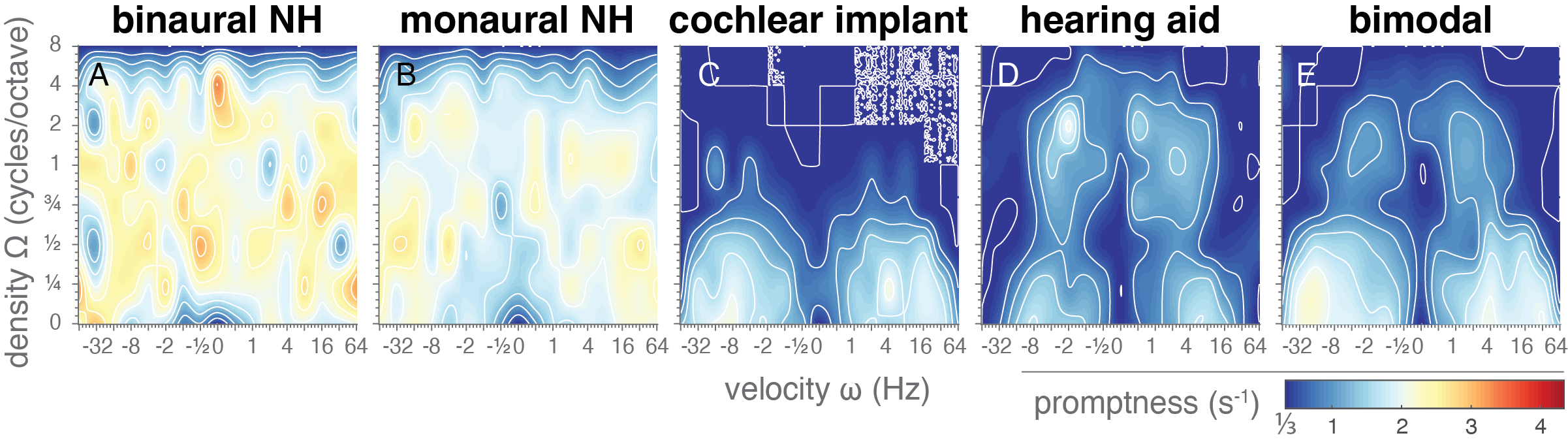

**Fig. S5. Spectrotemporal modulation transfer functions of listener L5.** Mean promptness as a function of velocity and density, representing spectrotemporal sensitivity in the normal hearing and simulated hearing-impaired conditions.

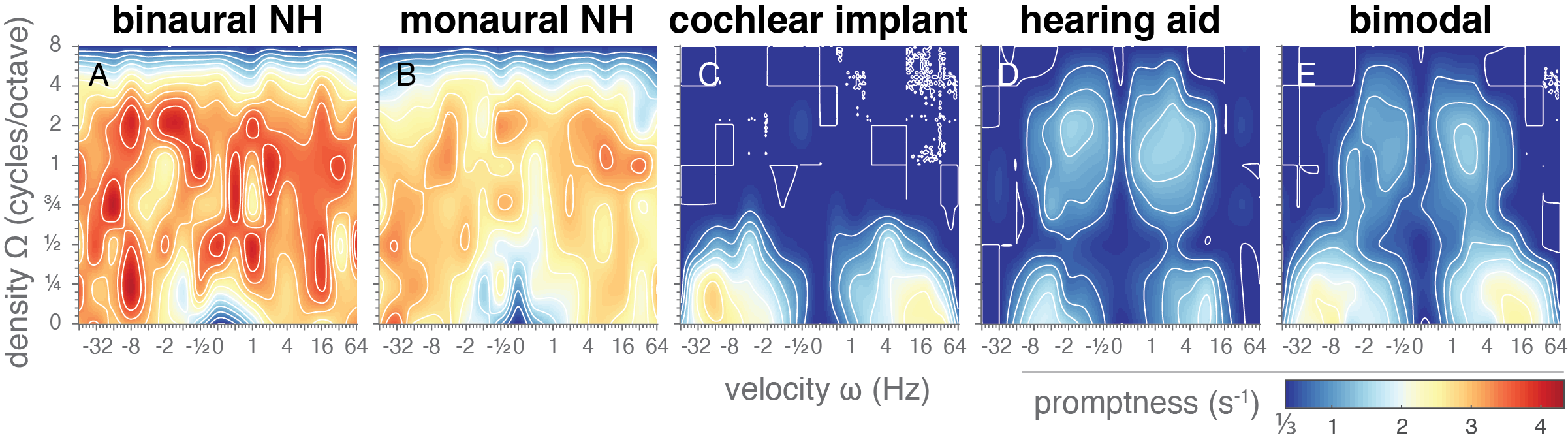

**Fig. S6. Spectrotemporal modulation transfer functions of listener L6.** Mean promptness as a function of velocity and density, representing spectrotemporal sensitivity in the normal hearing and simulated hearing-impaired conditions.

### Variability in the separability index

We assessed the degree of separability of the stMTF into a pure temporal and spectral component through singular value decomposition (SVD) using the separability index α_1_ (Eq. 5) and the α_2_ index (Eq. 7) to determine the relative contribution of the first (Eq. 4) and of the first two components (Eq. 6). In Fig. S7, these indices are determined for each listener separately, and the mean and 95% confidence interval are shown for each listening condition.

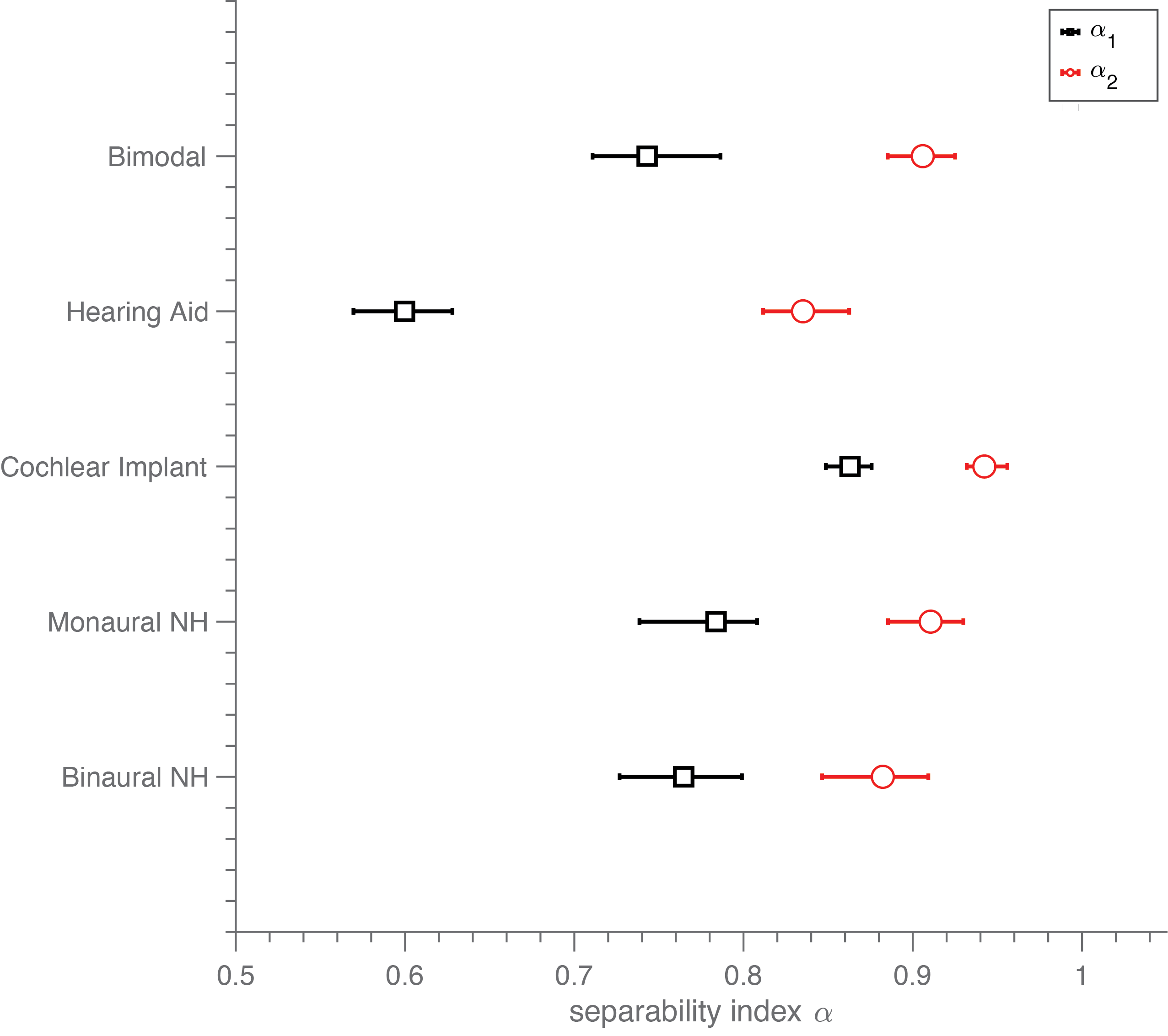

**Fig. S7. Separability index.** The separability index α_1_ (black squares) and the α_2_ index (red circles) are shown on the abscissa for each listening condition (ordinate). The squares and circles denote the mean estimate across listeners, while the vertical error bars denote the 95% confidence interval.
